# Evaluating hybridization capture with RAD probes as a tool for museum genomics with historical bird specimens

**DOI:** 10.1101/100867

**Authors:** Ethan B. Linck, Zach Hanna, Anna Sellas, John P. Dumbacher

## Abstract

Laboratory techniques for high-throughput sequencing have enhanced our ability to generate DNA sequence data from millions of natural history specimens collected prior to the molecular era, but remain poorly tested at shallower evolutionary time scales. Hybridization capture using restriction site associated DNA probes (hyRAD) is a recently developed method for population genomics with museum specimens (Suchan et al. 2016). The hyRAD method employs fragments produced in a restriction site associated double digestion as the basis for probes that capture orthologous loci in samples of interest. While promising in that it does not require a reference genome, hyRAD has yet to be applied across study systems in independent laboratories. Here we provide an independent assessment of the effectiveness of hyRAD on both fresh avian tissue and dried tissue from museum specimens up to 140 years old and investigate how variable quantities of input DNA affects sequencing, assembly, and population genetic inference. We present a modified bench protocol and bioinformatics pipeline, including three steps for detection and removal of microbial and mitochondrial DNA contaminants. We confirm that hyRAD is an effective tool for sampling thousands of orthologous SNPs from historic museum specimens to describe phylogeographic patterns. We find that modern DNA performs significantly better than historical DNA better during sequencing, but that assembly performance is largely equivalent. We also find that the quantity of input DNA predicts %GC content of assembled contiguous sequences, suggesting PCR bias. We caution against sampling schemes that include taxonomic or geographic autocorrelation across modern and historic samples.

## INTRODUCTION

Over the past three decades, novel laboratory techniques have enhanced our ability to generate DNA sequence data from millions of natural history specimens collected prior to the molecular era (Payne & Sorenson 2002). The advent of ancient DNA methods have allowed researchers to obtain both nuclear and mitochondrial DNA (mtDNA) sequences from extinct taxa (Cooper *et al.* 1992; Fleischer *et al.* 2006), explore changes in genetic diversity and population genetic structure over time (Weber *et al.* 2000; Habel *et al.* 2014), incorporate threatened or difficult-to-collect taxa into population genetic or phylogenetic studies (Guschanski *et al.* 2013; Linck *et al.* 2016), and take advantage of extant biological collections to boost sample size and inferential power (Wójcik *et al.* 2010; Linck *et al.* 2016). Now, high-throughput sequencing has dramatically increased both the overall efficiency of data collection and the total amount of sequence data that it is possible to collect from museum specimens (Rizzi *et al.* 2012; Hofreiter *et al.* 2015) by overcoming scalability hurdles intrinsic to traditional Sanger sequencing methods (Soltis & Soltis 1993; Wandeler *et al.* 2007).

Although high-throughput sequencing has already proved widely useful for incorporating museum specimens into phylogenomic studies (Burbano *et al.* 2010; McCormack *et al.* 2012; Besnard *et al.* 2015), its application for collecting genome-wide markers at the population level has lagged behind its use for addressing questions at deeper evolutionary time scales due to limitations in the most commonly employed library preparation methods for reduced-representation Illumina sequencing (Suchan *et al.* 2016). The limitations of historic museum samples include their high degree of fragmentation and low composition of long DNA fragments, which reduces the amount of flanking sequence that can be captured using ultraconserved element probes (Faircloth *et al.* 2012) and lowers the likelihood that multiple restriction digest recognition sequences are retained in a given DNA fragment (Baird *et al.* 2008; Peterson *et al.* 2012). Only in the past few years have library preparation protocols suitable for population genomics become available (Bi *et al.* 2013; Jones & Good 2016; McCormack *et al.* 2016), but their recent proliferation has meant that few have yet to be applied to multiple study systems in independent laboratories (McCormack *et al.* 2016). As a result, our understanding of the efficacy and biases of different approaches to reduced-representation genome sequencing from degraded DNA remains incomplete relative to either Sanger sequencing (Soltis & Soltis 1993; Wandeler *et al.* 2007) or high-coverage, single-sample whole genome sequencing (Poinar *et al.* 2006).

One promising, but under-tested approach to museum genomics suitable for population-level studies is hybridization capture of restriction site associated DNA (RAD) probes (hyRAD) (Suchan *et al.* 2016). Briefly summarized, the hyRAD method uses fragments produced by a double digest RAD (ddRAD) protocol (Peterson *et al.* 2012; Suchan *et al.* 2016) as the basis for biotinylated probes that capture orthologous loci in other samples, allowing them to be enriched and indexed for pooled Illumina sequencing. Although the method requires a high molecular weight DNA sample to produce the probe set, hyRAD offers advantages over other targeted capture methods in requiring no prior knowledge of the organism’s genome, such as transcriptome data or pre-existing sequences for probe design (Bi *et al.* 2013; McCormack *et al.* 2016). Additionally, because hyRAD relies on hybridization capture of orthologous regions across samples rather than retained restriction-site recognition sequences, the method mitigates the concerns of allelic dropout due to polymorphisms at restriction sites with increasing phylogenetic distance intrinsic to other RAD-based protocols (Gautier *et al.* 2013).

In their original paper, Suchan et al. (2016) validated their method by applying it to both fresh tissue and museum specimens of a butterfly *(Lycaena helle)* and grasshopper *(Oedaleus decorus).* They discussed the impact of library preparation, sample type, and bioinformatics pipeline on the number of SNPs produced. Here we provide an independent assessment of the effectiveness of hyRAD using both fresh avian tissues and dried tissue taken from museum specimens up to 140 years old. We present a modified version of the hyRAD protocol aimed at increasing efficiency and minimizing reagent use, and employ a custom bioinformatics pipeline with steps for detecting and removing microbial contamination in raw reads, contiguous sequences, and SNPs. We utilize hyRAD data to describe phylogeographic patterns in a New Guinea forest kingfisher *(Syma torotoro)* and we expand the available description of hyRAD’s performance by investigating how variable input DNA affects sequencing, assembly, and population genetic inferences.

## METHODS

### Study species, sampling, and DNA extraction

A major promise of museum genomics is the ability to conduct population-level studies in regions that are too logistically difficult to be amenable to broad modern sampling programs. The island of New Guinea is an apt example of this scenario, with poorly known biodiversity, large historical collections, rugged terrain, and ongoing political instability (Pratt and Beehler 2014; Dumbacher and Mack 2007). Phylogeographic research in New Guinea has been limited (Deiner et al. 2011; Dumbacher and Fleischer 2001), especially in the species inhabiting the island’s ring of lowland tropical rainforest. To evaluate the efficacy of for use in a broader study of the phylogeography of lowland new Guinea, we sampled 21 individuals of forest interior resident *Syma torotoro* (Yellow-billed Kingfisher), representing five named subspecies and the breadth of the species’ range on the island of New Guinea (**Table 1**). For seven individuals, we extracted whole-genomic DNA from fresh tissue using a DNeasy tissue extraction kit (Qiagen, Valencia, CA) following the manufacturer’s protocol. For the remaining 14 individuals, we extracted DNA from the toe-pads of museum study skins in a dedicated ancient DNA laboratory at the California Academy of Sciences using a phenol-chloroform and centrifugal dialysis method. No modern DNA or post-PCR products are handled in this lab, which is located on a separate floor from the main genetics facility.

**Table 1.** Sampling information.

| Specimen | Subspecies | Sample Type | Locality | Date collected | Age of sample (years) | Total number of reads | Number of cleaned reads | % Duplicate reads | Average Coverage (X) | Specificity | Fold Enrichment | Sensitivity | Loci | Average locus length (nt) | Number of contigs over 1KB | Initial concentration (ng/μL) | GC content of on-target contigs |
| --- | --- | --- | --- | --- | --- | --- | --- | --- | --- | --- | --- | --- | --- | --- | --- | --- | --- |
| KU:Birds:5215 | *pseustes* | Tissue | Ivinka Camp, Gulf Province, Papua New Guinea | 2003 | 13 | 18,713,496 | 2,915,214 | 68.1 | 12.74 | 16.87 | 16.58 | 84.93 | 22,568 | 518 | 489 | 71.6 | 44.22 |
| KU:Birds:5464 | *pseustes* | Tissue | Sapoa Camp, Gulf Province, Papua New Guinea | 2003 | 13 | 17,246,396 | 2,512,928 | 68.2 | 11.2 | 16.65 | 16.36 | 79.43 | 19,880 | 499 | 340 | 51.8 | 44.69 |
| KU:Birds:6927 | *meeki* | Tissue | Mt. Suckling, Oro Province, Papua New Guinea | 2011 | 5 | 25,892,086 | 3,103,912 | 73.2 | 14.46 | 17.37 | 17.07 | 85.02 | 23,162 | 510 | 494 | 37 | 44.53 |
| KU:Birds:7131 | *pseustes* | Tissue | Dark End Camp, Gulf Province, Papua New Guinea | 2002 | 14 | 6,819,041 | 1,154,541 | 60.4 | 5.87 | 17.29 | 16.99 | 65.44 | 12,725 | 434 | 157 | 31.8 | 45.01 |
| CAS:ORN:626 | *meeki* | Blood | Varirata National Park, Central Province, Papua New Guinea | 2011 | 5 | 15,548,547 | 1,945,868 | 70.2 | 9.38 | 17.06 | 16.77 | 77.25 | 18,519 | 488 | 262 | 43 | 44.62 |
| KU:Birds:9145 | *torotoro* | Tissue | Gahom Camp, East Sepik Province, PNG | 2003 | 13 | 18,049,058 | 2,309,716 | 70.2 | 10.75 | 16.91 | 16.62 | 80.15 | 20,057 | 452 | 312 | 10.1 | 44.35 |
| AMNH:Birds:637445 | *torotoro* | Toe pad | Wasior, West Papua Province, Indonesia | 1928 | 88 | 13,112,780 | 48,257 | 78.2 | 2.59 | 11.16 | 10.97 | 33.52 | 849 | 449 | 57 | 3.66 | 46.03 |
| AMNH:Birds:329542 | *ochracea* | Toe pad | Sewa Bay, Normanby Island, Papua New Guinea | 1934 | 82 | 15,875,059 | 40,402 | 82.2 | 1.8 | 8.9 | 8.75 | 29.19 | 388 | 465 | 24 | 3.84 | 47.62 |
| AMNH:Birds:637464 | *tentelare* | Toe pad | Wannambai, Maluku Province, Indonesia | 1896 | 120 | 21,058,924 | 139,248 | 59.9 | 9.2 | 11.07 | 10.88 | 50.24 | 3,197 | 449 | 154 | 10.2 | 44.98 |
| AMNH:Birds:637450 | *torotoro* | Toe pad | Humbolt Bay, Papua Province, Indonesia | 1928 | 88 | 3,692,636 | 16,359 | 75.1 | 0.89 | 12.06 | 11.85 | 14.62 | 153 | 543 | 15 | 0.714 | 48.74 |
| AMNH:Birds:637429 | *torotoro* | Toe pad | Misool Island, West Papua Province, Indonesia | 1900 | 116 | 13,531,142 | 35,925 | 79.8 | 2.13 | 9.46 | 9.30 | 30.09 | 412 | 448 | 26 | 2.54 | 46.44 |
| AMNH:Birds:300723 | *torotoro* | Toe pad | Waigeu Island, West Papua Province, Indonesia | 1900 | 116 | 14,143,466 | 34,596 | 82.2 | 1.91 | 10.03 | 9.86 | 27.78 | 508 | 401 | 21 | 0.846 | 46.88 |
| AMNH:Birds:637446 | *torotoro* | Toe pad | Kepaur, West Papua Province, Indonesia | 1897 | 119 | 20,487,169 | 36,627 | 84.1 | 2.15 | 8.02 | 7.88 | 23.19 | 177 | 600 | 22 | 1.07 | 48.18 |
| NHMK:ZOO:1911.12.20.823 | *pseustes* | Toe pad | Mimika River, Papua Province, Indonesia | 1913 | 103 | 2,875,704 | 19,115 | 50.9 | 1.55 | 10.85 | 10.66 | 20.44 | 239 | 611 | 40 | 1.49 | 47.28 |
| NHMK:ZOO:1911.12.20.822 | *pseustes* | Toe pad | Satakwa River, Papua Province, Indonesia | 1911 | 105 | 5,464,605 | 46,548 | 59.8 | 2.08 | 9.44 | 9.28 | 24.39 | 296 | 579 | 36 | 0.584 | 48.32 |
| AMNH:Birds:437798 | *torotoro* | Toe pad | Amberbaki, West Papua Province, Indonesia | 1877 | 139 | 1,579,610 | 4,957 | 73.4 | 0.39 | 9.1 | 8.94 | 4.58 | 52 | 582 | 4 | 1.84 | 48.88 |
| AMNH:Birds:637441 | *torotoro* | Toe pad | Mt. Mori, West Papua Province, Indonesia | 1899 | 117 | 6,803,386 | 19,623 | 76.9 | 1.3 | 9.5 | 9.34 | 21.38 | 157 | 512 | 11 | 7.36 | 47.89 |
| AMNH:Birds:293741 | *torotoro* | Toe pad | Ifaar, Papua Province, Indonesia | 1928 | 88 | 27,602,106 | 49,786 | 91.1 | 2.14 | 12.48 | 12.27 | 23.84 | 493 | 447 | 28 | 0.124 | 50.41 |
| AMNH:Birds:293715 | *torotoro* | Toe pad | Kepaur, West Papua Province, Indonesia | 1928 | 88 | 30,751,249 | 56,972 | 90.5 | 2.15 | 11.15 | 10.96 | 24.2 | 438 | 504 | 30 | 0.15 | 50.95 |

### Library Preparation, Hybridization Capture Experiments, and Sequencing

We prepared samples for reduced-representation whole-genome sequencing using a modified version (Hanna & Sellas 2016) of Suchan *et al*’s hyRAD method (Suchan *et al.* 2016) aimed at increasing efficiency of reactions and reducing reagent use. We present this protocol in a detailed bench-ready version online (https://github.com/calacademy-research/hyRADccg), and summarize it below.

To produce biotinylated probes, we performed a double restriction digest with enzymes MluCl and SphI (New England Biolabs) on 400 ngs of high molecular weight DNA extracted from fresh tissue of a single *S. t. ochracea* individual. After ligation of adapters to fragments, we size-selected the resulting fragments on a Pippin Prep (Sage Science, Beverly, USA) with a target peak at 270 bp and ‘tight’ size selection range. We ran 16 cycles of real-time polymerase chain reaction (RT-PCR), and purified products by gel excision and a Zymoclean Gel Recovery Kit (Zymo Research). We preserved one aliquot of this product for sequencing while performing an additional MluCl / SphI double digest to de-adapterize a second aliquot. We labeled this deadapterized aliquot with biotin-14-dATP, using a BioNick DNA Labeling System (Thermofisher Scientific).

To produce whole genome libraries, we sheared high molecular weight DNA from modern tissue samples to ˜400 bps using a M-220 Focused-ultrasonicator (Covaris). DNA from museum specimen toepads was already fragmented as a product of natural degradation associated with the age of the samples, and was therefore left untreated. For both modern and historic samples, we used a Kapa Hyper Prep Kit (Kapa Biosystems) to prepare dual-indexed libraries. We amplified libraries using 5–13 cycles of RT-PCR. After quantifying DNA content in each sample, we made standardized dilutions of each sample and combined equal amounts of these dilutions to create one pool of modern DNA samples (n=6) and two pools of ancient DNA samples (n=7 each). We used a 1-1.5x ratio of of AmPure XP beads to remove small DNA fragments throughout the protocol, and assessed DNA quantity and quality with a Qubit 2.0 fluorometer and an Agilent 2100 Bioanalyzer between all major steps.

To perform hybridization capture reactions, we incubated each pool of samples with 250 ng of biotinylated probe for 48-72 hours at 55°C in a solution containing 20X SSC, 50X Denhardt’s Solution, 0.5M EDTA, 10% Sodium Dodecyl Sulfate (SDS), and a blockers mix containing Chicken Hybloc (0.5 μg/μL), IDT’s xGen Universal Blocking Oligo -TSHT- i5 (0.05 nmole/μL), and IDT’s xGen Universal Blocking Oligo -TSHT- i7 (0.05 nmole/μL). Following hybridization, we prepared 50 μL Dynabeads MyOne Streptavidin C1 beads for use by washing three times with 1X binding buffer containing 2 M NaCl, 10 mM Tris-HCl (pH 7.5), 0.5% Polysorbate 20 (Tween 20), and 1 mM EDTA, and final resuspension in 70 μL 2X binding buffer. We then bound the probes to the beads by mixing and incubating at room temp for 30 min. After performing three 500 μL washes of the bead-probe mixture using a pre-warmed buffer containing 10% Sodium Dodecyl Sulfate (SDS) with 0.1X saline-sodium citrate (SSC), 0.1% SDS buffer, we concentrated our final pooled libraries in 30μL 10 mM Tris-HCl, 0.05% Tween 20 (pH 8-8.5). We next amplified these libraries using RT-PCR for 9-12 cycles, cleaned using 1.2X Ampure XP beads, and quantified using Qubit. We sent a single final pool with equimolar amounts of all three hybridized pools to University of California Bekeley’s QB3 Vincent J. Coates Genomics Sequencing Laboratory (hereafter called “QB3”) for sequencing with 100 bp paired-end sequence reads on a single lane of an Illumina HiSeq 4000.

### Sequence read quality control, assembly, and alignment

To clean and quality filter reads, assemble reads into contigs, align sequences across samples, and map reads to merged alignments for SNP discovery, we used a custom pipeline combining in-house R scripts as well as preexisting genomics tools and wrapper scripts from QB3’s two *de novo* targeted capture bioinformatics pipelines (https://github.com/CGRL-QB3-UCBerkeley; “denovoTargetCapturePopGen” and “denovoTargetCapturePhylogenomics”). We present our full pipeline online as both a tutorial and a list of shell commands (https://github.com/elinck/hyRAD/) **(Figure 1).**

**Figure 1.**
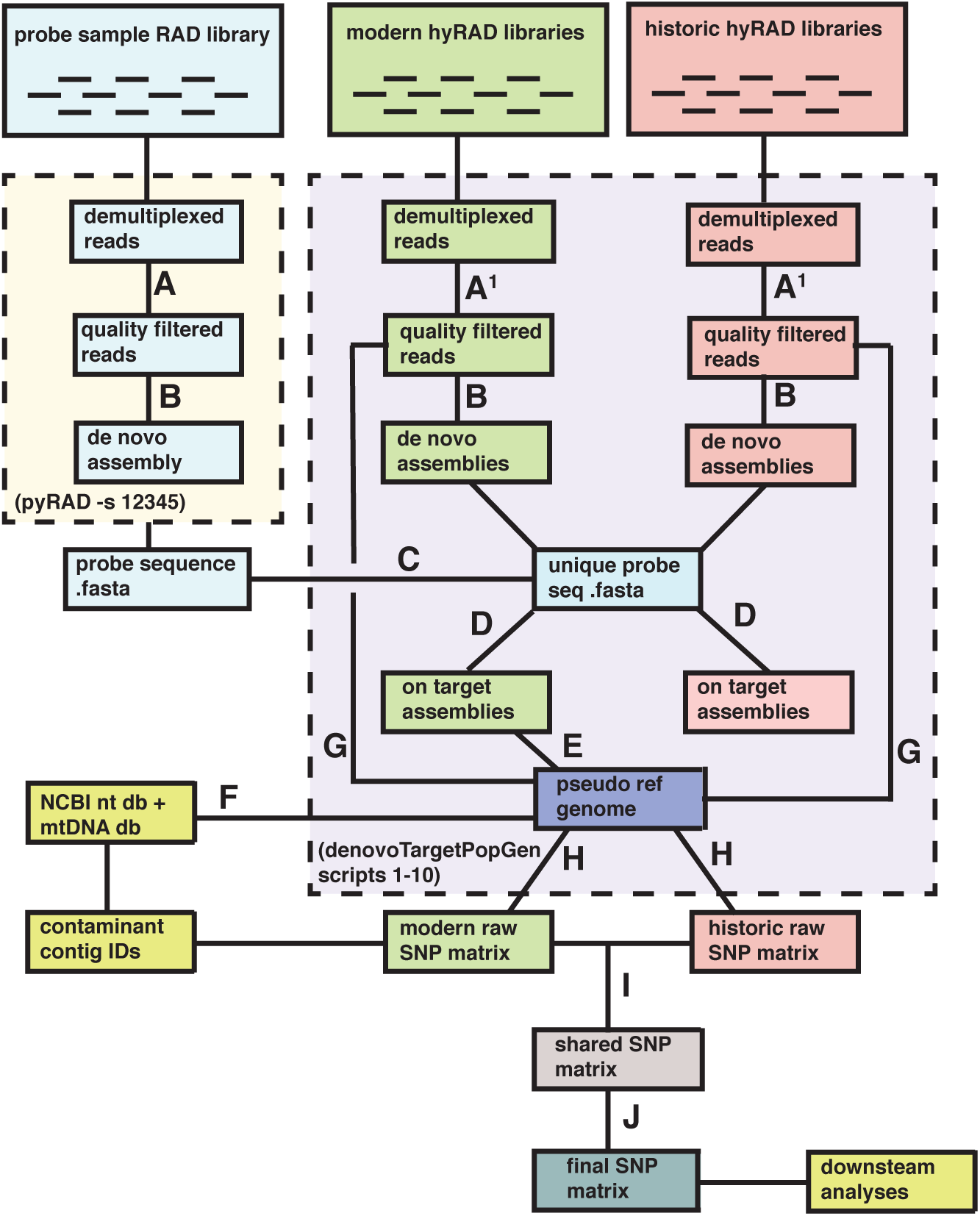
Bioinformatics pipeline for *S. torotoro* hyRAD data. We demultiplexed 100 bp paired- end reads from three genomic libraries and filtered for adapter contamination / quality scores **(A)**, or adapter contamination / quality scores and E. coli contamination (A1). Reads were clustered (as consensus fasta files) **(B)**, and repeat regions removed from probes **(C)**. After determining which assembled clusters were orthologous with probe regions **(D)**, we merged flanking regions from on-target loci in modern samples with the repeat-free probe sequence to create a pseudo-reference genome **(E)**. To identify which contigs represented contamination in the original probe sample library from exogenous microbes or mitochondrial DNA, we BLAST searched against both the NCBI nt database and a full mitochondrial genome from S. torotoro relative Halcyon sanctus **(F)**. We aligned quality filtered reads to this pseudo reference **(G)**, called SNPs to produce a raw .vcf file for historic and modern DNA libaries separately **(H)**. After filtering SNPs for origin in contaminant contigs and then restricting our matrix to sites present in both sample types **(I)**, we filtered SNPs by read depth, quality scores, probability of being variable sites, and minor allele frequencies **(J)** prior to downstream analyses.

We first processed reads from our probe library with pyRAD version 2.17 (Eaton 2014) to create a pseudo reference genome to use as the basis for aligning sequences from samples in our hybridization capture reactions. After quality-filtering reads and trimming adapter contamination, pyRAD used the vsearch algorithm (Rognes *et al.* 2016) to cluster reads into loci within samples, cluster loci into stacks between samples, and aligned putatively orthologous loci using MUSCLE version 3.8 (Edgar 2004). We implemented strict adapter filtering, retained reads longer than 70 bp after trimming, set a minimum sequence identity threshold of 97% for clustering, and kept four sites per cluster with a Phred Quality Score <20. We removed repetitive genomic regions with the NCBI BLAST+ version 2.4 tool BLASTn (Boratyn *et al.* 2013) by aligning the output file of assembled clusters against itself, and retained only cluster sequences that aligned uniquely to themselves using an e-value of 0.00001.

To remove reads that failed to pass Illumina quality control filters, trim reads for quality and adapter contamination, merge overlapping reads, remove PCR duplicates, and remove endogenous *E. coli* contamination, we used QB3’s denovoTargetCapturePopGen “2-ScrubReads” wrapper around the Trimmomatic version 0.36 (Bolger *et al.* 2014), Bowtie 2 (Langmead 2010), Cutadapt (Martin 2011), Cope (Liu *et al.* 2012), FastQC (http://www.bioinformatics.babraham.ac.uk/projects/fastqc/), and FLASh (Magoc & Salzberg 2011) tools. We assembled cleaned and filtered reads for each sample using QB3’s denovoTargetCapturePhylogenomics wrapper script “2-GenerateAssembliesPhylo” around the SPAdes version 3.8.1 genome assembler (Bankevich *et al.* 2012), which automatically selects a k-mer value based on read length and dataset type. To determine which contigs from our capture libraries were orthologous with probe regions, we used the denovoTargetCapturePopGen wrapper “5-FindingTargets” around the BLAST+ (Boratyn *et al.* 2013) and cd-hit-est (Fu *et al.* 2012) tools. Analyzing samples from modern and historical DNA separately, we used a clustering identity threshold of 95% and permitted 100 bp of sequencing flanking the core probe region. After determining matches, we collapsed overlapping, orthologous contigs from all modern samples with the probe library to generate an extended pseudo-reference genome to which we aligned cleaned reads using QB3’s denovoTargetCapturePopGen wrapper “7- Alignment” around the Novoalign version 3.04.06 tool (http://www.novocraft.com/products/novoalign/). We ran Novoalign with an average library insert size of 235 and a maximum alignment score of 90.

### SNP discovery

Traditional SNP calling algorithms based on allele counting and quality scores are characterized by high degrees of uncertainty with low-coverage sequence data (Korneliussen *et al.* 2014). We incorporated uncertainty into genotype estimation by calling SNPs and estimating allele frequencies using an empirical Bayesian framework implemented in the software ANGSD version 0.913 (http://www.popgen.dk/angsd/index.php/ANGSD). ANGSD uses the likelihood of all ten possible genotypic configurations for each site passing quality filters in all individuals to estimate a site frequency spectrum (SFS), which is then used as a prior to estimate the posterior probabilities for all possible allele frequencies at each site in each sample. Using these estimates, we called SNPs with a 95% probability of being variable and a minimum minor allele frequency of 5%.

### Contamination control and data filtering

In order to identify if any contigs in our assemblies represented off-target mtDNA captures, we performed a BLAST+ (Boratyn *et al.* 2013) nucleotide search with each of our assemblies as a query against a database of the full mitochondrial genome of *S. torotoro* relative *Halcyon santcus*. We then removed all matching contigs from each sample’s assembly fasta with in-house R scripts (“excerptcontigIDs.R” and “cutcontigsbatch.R”), and used these mtDNA-free sequences for all subsequent assembly performance calculations. To prepare our sequence alignment in .sam/.bam format for SNP calling, we followed Bi et al. (2013) in hierarchically filtering out individuals, contigs, and sites that appeared to be quality outliers, and implemented additional steps for regions derived from microbial contamination or mitochondrial DNA. We determined no individuals had abnormal coverage (defined as less than 1/3 or greater than 3X the average coverage across all individuals) and we created merged, sorted BAM files and generated raw variant call format files (.vcf) with samtools version 1.3 (Li *et al.* 2009) and bcftools version 1.3.1 (Narasimhan *et al.* 2016), processing modern and historical DNA samples separately.

Because our SNP matrix was derived independently from our assemblies, we performed a second step of contamination filtering by removing SNPs originating from read alignments to regions of exogenous microbial DNA and / or mtDNA present in the original probe sample RAD library. We used our full pseudo-reference genome as the query in a search of the entire BLAST+ (Boratyn *et al.* 2013) nucleotide database and our *H. sanctus* mtDNA BLAST+ database. We then used Henderson *et al.’s* (2016) “GItaxidIsVert.py” script to identify sequences that were potentially microbial in origin, and performed a second BLAST+ search with this subset to further select only the subset of contigs that had their best or only alignment with nonvertebrate reference genomes. To exclude such sequences as well as those aligning with mitochondrial DNA sequence, we used the vcftools version 0.1.11 “--not-chr” flag, and removed indels in the same step with the “--remove-indels” flag.

We estimated independent empirical gene coverage and site depth distributions using QB3’s denovoTargetCapturePopGen “9-preFiltering” script and used these distributions as input to the QB3 “10-SNPcleaner” script. Run separately for modern and historical samples, this script removed all sites with coverage below 6x, sites missing in more than half of our samples, sites with biases associated with quality score, mapping quality, or distance of alleles from the ends of reads. Because hydrolytic deamination of cytosine (C) to uracil (U) residues is the most common form postmortem nucleotide damage present in historic museum specimens, which may result in misincorporation of thymines (Ts) instead of uracil during PCR amplification and bias population genetic inference, this script also eliminated all C to T and G to A SNPs (Hofreiter *et al.* 2001; Briggs *et al.* 2007; Axelsson *et al.* 2008). Finally, we used the BEDtools version 2.26.0 “intersect” function (Quinlan & Hall 2010) to retain only the sites that passed all filters for both historic and modern specimens.

### Statistical analyses

To assess the differences in sequencing and assembly performance between modern and historical samples, we implemented Wright’s Two-Sample T-Tests in the R (R Core Team 2016). We evaluated differences between groups in the mean percentage of duplicate reads, the mean number of on-target contigs, the mean length of on-target contigs, and the percentage of sequenced reads successfully mapping to our pseudo-reference genome. Because some commonly used polymerases bias against amplification of targeted DNA in favor of the GC-rich microbial contamination common to extracts from museum specimens (Dabney & Meyer 2012), we assessed differences in GC content present in assembled on-target contigs. In order to determine whether sample age, initial sample DNA concentration, or sequencing effort were significant predictors of %GC content among historical samples and mean number or length of on-target contigs, we used simple linear regression. We used stepwise model selection with corrected Akaike Information Criterion (AICc) scores to determine best-fit models, and did not include interaction terms to avoid over-parameterization given our small sample size.

### Population Genetic Clustering and Discriminant Analysis of Principal Components

Although accurate estimation of population genetic structure in *Syma torotoro* was not the primary goal of our study, we were nonetheless interested in assessing hyRAD’s ability to produce biologically meaningful results by testing whether our data reflected the signature of phylogeographic processes such as isolation by distance and vicariance, rather than the signature of DNA degradation, contamination, or other artifacts of library preparation and sequencing. We implemented k-means clustering and Discriminant Analysis of Principal Components (DAPC) in the R package adegenet (Jombart 2008), using 100% complete data matrix (1,690 SNPs) to avoid biasing inferences with nonrandom patterns of missing data. We retained all principal component (PC) axes for k-means clustering and inspected both population assignments and change in Bayesian Information Criterion (BIC) scores across multiple values of K to select an optimal partitioning scheme. To maximize among-population variation and calculate ancestral population membership probabilities for each sample, we performed DAPC on the first six PCs using two discriminant axes. We then chose to retain these six PCs to optimize the a*-*score value for our data, which is the difference between the proportion of successful reassignment of the analysis and values obtained using random groups. However, because the change in BIC scores failed to clearly indicate any “true” value of K, we repeated our analysis with clustering assignments for K values of 1-8. To explore correlations between our three retained PCs and variables expected to differ between modern and historic samples (specificity, sensitivity, fold enrichment, age, initial concentration), we again performed simple linear regressions. Finally, to test for patterns of isolation by distance (IBD) across our samples, we performed a Mantel test among all individuals based on 999 simulated replicates using the R package ade4 (Thioulouse & Dray 2007).

## RESULTS

### Hybridization Capture Experiments and Sequencing

We obtained a total of 397 million sequence reads for the probe and hybridization capture libraries, successfully demultiplexing 20/21 samples, with one sample failing due to bar code error. The total reads per sample ranged from 1.6 million to 30.7 million, and the average number of reads per sample did not vary significantly between modern and historical samples (t=-0.946, df=14.6, p=0.359). Of the original 397 million reads, 19.8% passed initial Illumina quality filters, contamination checks, adapter trimming, and removal of PCR duplicates **(Table 1).** The resulting number of cleaned reads per sample ranged from approximately 16,000 to 3.1 million, with an average count of 766,662, and significantly fewer reads for historic samples (t=-3.185, df=13.088, p=0.007). The majority of reads lost to quality control were PCR duplicates, with a range of 50.9% to 91.1% duplicate reads per sample. 2,455 reads were removed as *E. coli* contamination from 12 of 20 individuals (range 1 to 2,318 reads per individual) **(Table 2).** The average depth of read coverage per sample, calculated as the read depth per base averaged across the length of the pseudo reference genome, ranged from 7.6 to 26.4, and was significantly lower in historic samples (*t*=- 3.754, df=12.632, *p*=0.002)

**Table 2.**
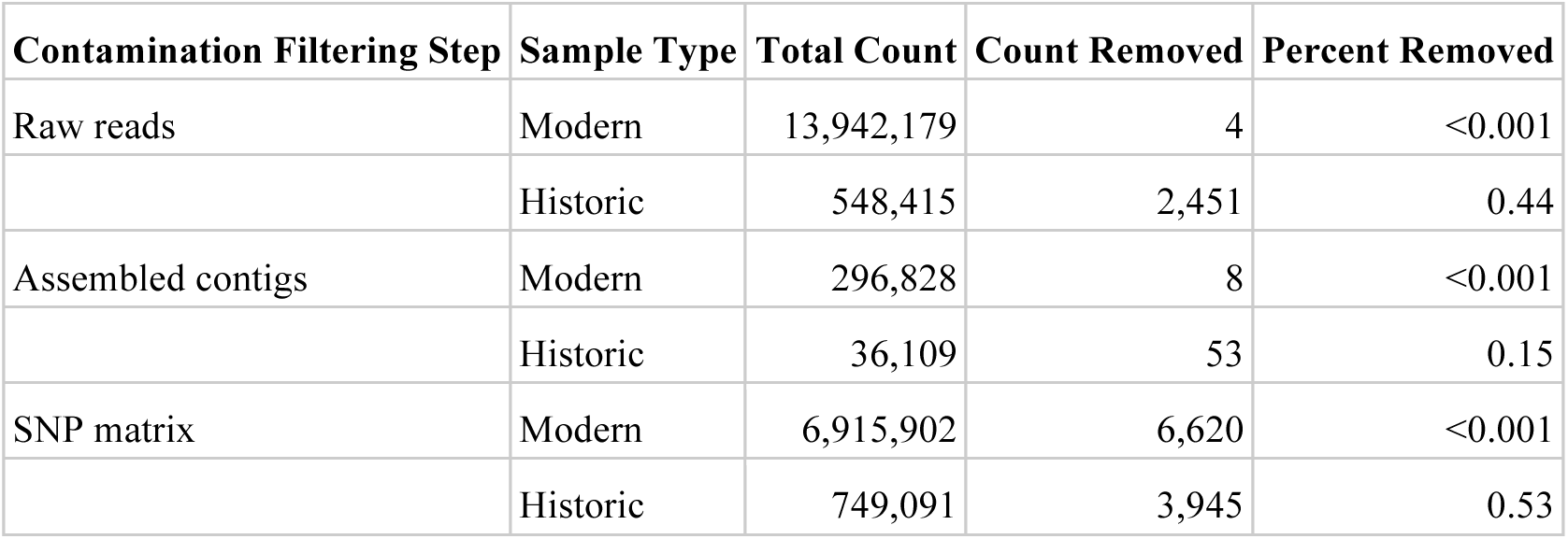
Results of microbial and mitochondrial contamination removal at three distinct steps. Raw reads were filtered for *E. coli* contamination; assemblied contigs were filtered for mitochondrial DNA; SNP matrices were filtered for both microbial DNA and mitochondrial DNA.

| Contamination Filtering Step | Sample Type | Total Count | Count Removed | Percent Removed |
| --- | --- | --- | --- | --- |
| Raw reads | Modern | 13,942,179 | 4 | <0.001 |
|  | Historic | 548,415 | 2,451 | 0.44 |
| Assembled contigs | Modern | 296,828 | 8 | <0.001 |
|  | Historic | 36,109 | 53 | 0.15 |
| SNP matrix | Modern | 6,915,902 | 6,620 | <0.001 |
|  | Historic | 749,091 | 3,945 | 0.53 |

### Assembly and alignment results

Assembly of our probe library resulted in a total of 687,318 contigs and 61.9 million nucleotides (nt), which was reduced to 502,820 unique contigs and 16.1 million nt after excluding repetitive regions. We captured orthologous loci from all successfully sequenced samples. The number of on-target contigs per sample ranged from 55 to 23,155, with a significantly higher average number of on-target contigs among modern samples **(Table 1; Figure 2)**. Among our assemblies, we removed eight contigs from the modern samples and 53 contigs from the historic samples due to their mitochondrial origin **(Table 2)**. Mean contig size ranged from 401–611 nt, and did not differ significantly between modern and historic samples. However, the historic samples had significantly fewer contigs exceeding 1 knt in length (t=–3.181, df=5.004, p=0.025). The percentage of reads passing quality filters that successfully mapped to the pseudo-reference genome (also known as specificity) ranged from 51.8-57.7%, and was significantly higher on average for modern samples **(Figure 2).** Additionally, %GC content was significantly higher in historic than modern samples **(Figure 2).** Among historic samples, the number of cleaned reads was a significant predictor of the number of captured loci and sample input DNA quantity was a significant predictor of %GC content in on-target assembled contigs **(Figure 3).**

**Figure 2.**
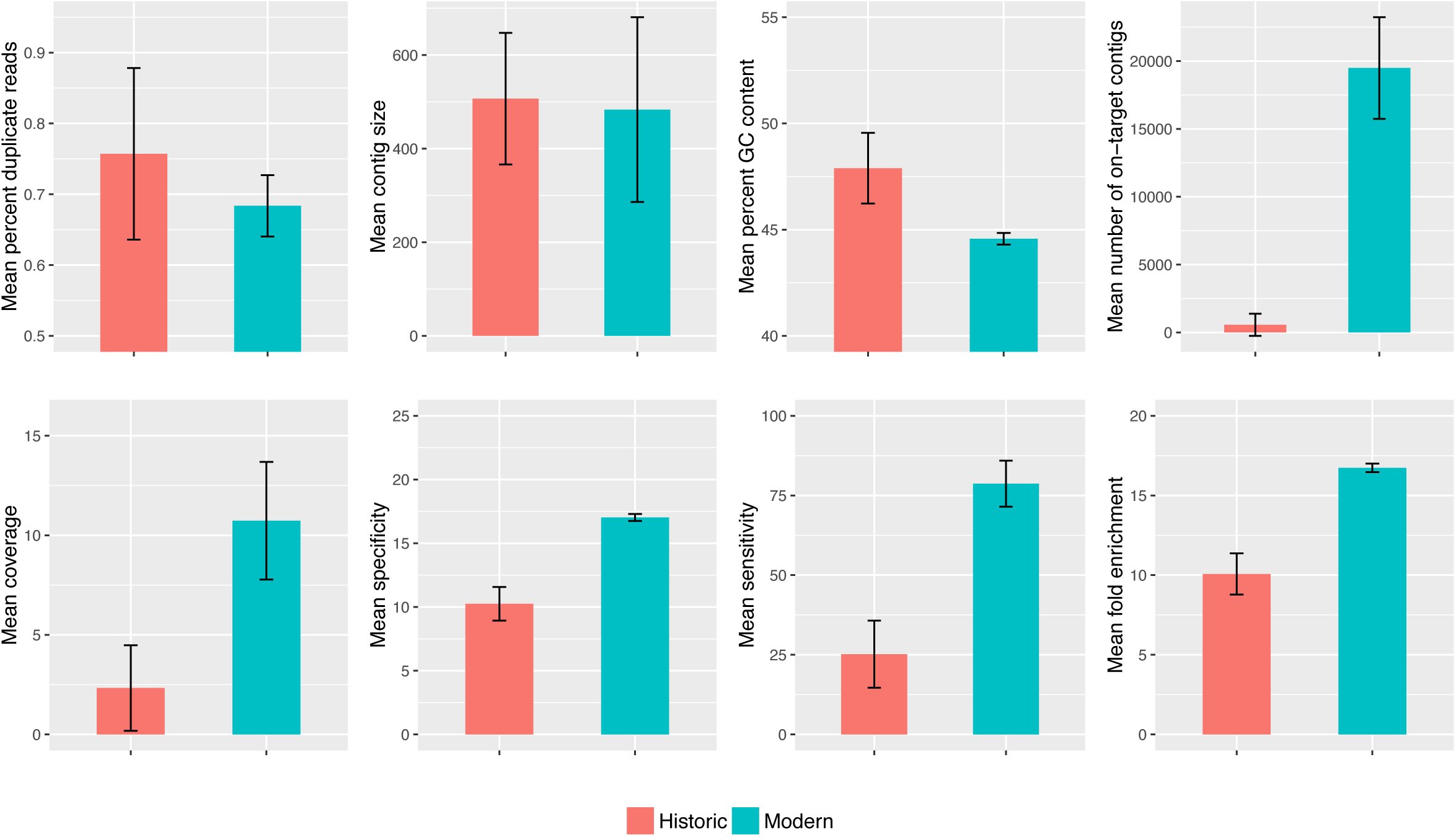
Differences in sequencing and assembly performance between historical and modern DNA extractions. We observed significantly higher specificity (×=−17.711, df=14.015, *p*<0.001), sensitivity (×=−12.928, df=14.014, *p*<0.001), fold enrichment (×=−17.711, df=14.015, *p*<0.001), and average coverage (×=−6.248, df=7.555, *p*<0.001) in modern samples. We recovered a significantly higher total number of on-target loci in modern samples (×=−12.239, df=5.2211, p<0.001), but significantly higher mean percent GC content in historic samples (×=6.997, df=13.368, *p*<0.001). We observed no significant differences between modern and historic samples for mean contig size or mean percentage of duplicate reads.

**Figure 3.**
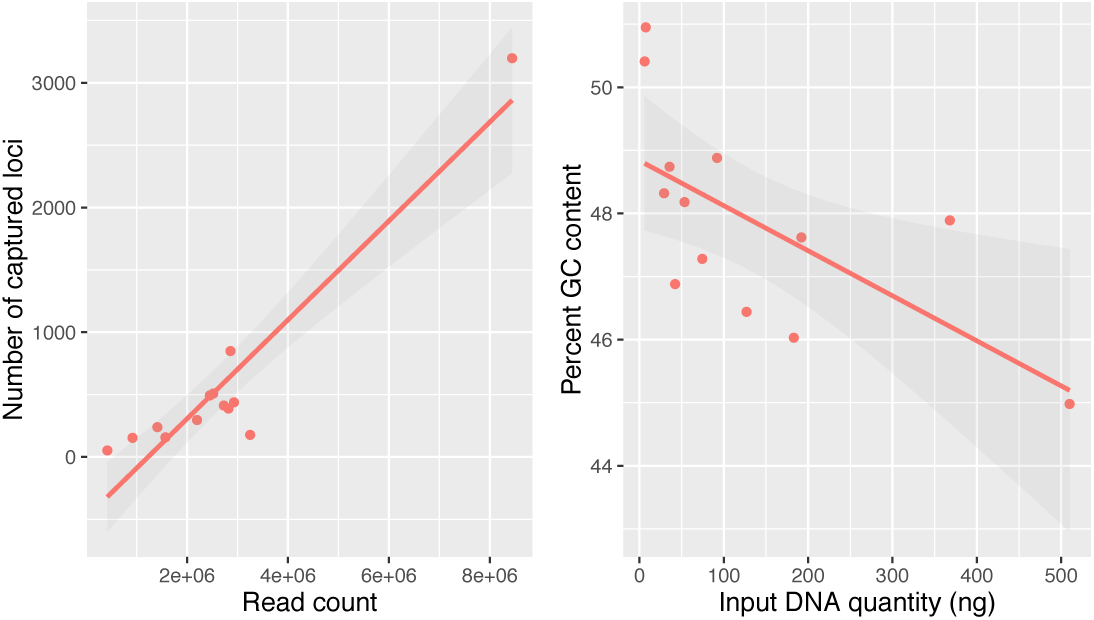
Among historic samples, the number of trimmed reads was a significant predictor of the number of captured loci (R^2^=0.872, p=1.925e-06) and the initial sample DNA input quantity was a significant predictor of percent GC content in on-target assembled contigs (R^2^=0.370, *p*=0.016).

### SNP discovery and filtering

We identified two contigs of mitochondrial origin and eight contigs of potential nonvertebrate origin in our pseudo-reference genome, and excluded all sites from these contigs in our alignment prior to SNP calling **(Table 2)**. Using ANGSD, we identified 39,105 high quality SNPs with at least a 95% probability of being variable, with a matrix completeness of 62.8% (or 37.2% missing data across all individuals) and an average of 12.2 SNPs per assembled contig. Per individual, the proportion of missing sites ranged from 5.6% to 90.6%, with a significantly higher mean percentage missing data for historic samples (54.5%) than modern samples (28.3%) (t=6.727, df=14.594, p=7.812e-06). The total number of SNPs in our data matrix decreased linearly after the first 10 individuals when we increased the minimum number of individuals successfully genotyped in order to retain each SNP **(Figure S1).**

### Population Genetic Clustering and Discriminant Analysis of Principal Components

DAPC analysis of our 100% complete data matrix revealed a linear pattern of increase in the total amount of genetic variation explained when retaining additional principal components. Replicate attempts to optimize a-score values alternatively suggested retaining either five or six PCs to maximize discrimination ability without overfitting the model. Similarly, BIC scores from DAPC decreased in an approximately linear fashion as more clusters were added and did not indicate a clear shift to a slower rate of BIC change. Therefore, we repeated our analysis for values of K from 1-8, which revealed patterns of increasingly fine, geographically coherent structure from K=1 to K=5 **(Figure 5).** At K=2, DAPC separated individuals from mainland Papua New Guinea (PNG) from individuals in western New Guinea and Normanby Island in PNG, which also reflected the break between modern and historic samples. At K=3, DAPC identified an additional cluster from West Papua that included individuals from the southwest New Guinea Coast and the Aru Islands. At K=4, DAPC isolated an individual from the northern slope of the Arfak Mountains in Western New Guinea, which, collected in 1877, was also the oldest sample included in our study. A fifth cluster distinguished the single individual from Normanby Island, PNG. At K=6 and greater, DAPC began subdividing individuals into additional clusters without a shared geographic basis. Our Mantel test did not find statistically significant correlation between geographic and genetic distance (p=0.251). Linear regression analyses showed significant correlations between all variables and PC1 but no other PC and sample variable pairs **(Table S1).**

## DISCUSSION

### Hybridization-RAD is an effective tool for sampling thousands of orthologous SNPs from historic museum specimens

In their description of hyRAD, Suchan et al. (2016) suggest that it allows “sequencing of orthologous loci even from highly degraded DNA samples” and can be used to retrieve sequence data using museum samples up to 100 years old.

Our hybridization capture experiments in an independent laboratory with an independent study organism *(Syma torotoro)* largely support this conclusion. Our modified protocol had comparable or better performance to the Suchan *et al.* (2016) protocol applied to the butterfly *Lycaena helle* in terms of the number of assembled contigs, orthologous loci, total SNPs collected, and number of SNPs recovered across multiple individuals **(Table 1; Figure 4).** In particular, we note that our application of identical library preparation methods for both modern and historic libraries appears to have been at least equally effective to the Suchan *et al.* (2016) method, potentially mitigating the need for separate protocols by sample type. While much of the perceived improvement could be attributed to an increased number of probes resulting from the number of restriction site recognition sequences scaling with genome size (˜269 MB for *L. helle* and ˜1.5 GB for Gregory 2017), it also reflects the robustness of the approach to different taxa, laboratory conditions, specimen preparation conditions, and bioinformatics pipelines. Even with stringent filtering for quality and post-mortem damage, our 100% complete data matrix of 1,690 SNPs is comparable to or exceeds the number of orthologous SNPs collected in similar studies of museum genomics that used UCE capture (McCormack *et al.* 2016), exon sequence capture (Bi et al. 2013), or even ddRAD methods with fresh tissues (Shultz *et al.* 2016). For phylogenetic or population genetic analyses methods that correct for nonrandom patterns of missing data, our full matrix of 39,105 SNPs potentially offers significant power to resolve rapid, recent divergences, detect fine scale patterns of population structure, infer historical effective population sizes with high accuracy, and reveal histories of drift, selection, and migration (Toews *et al.* 2016).

**Figure 4.**
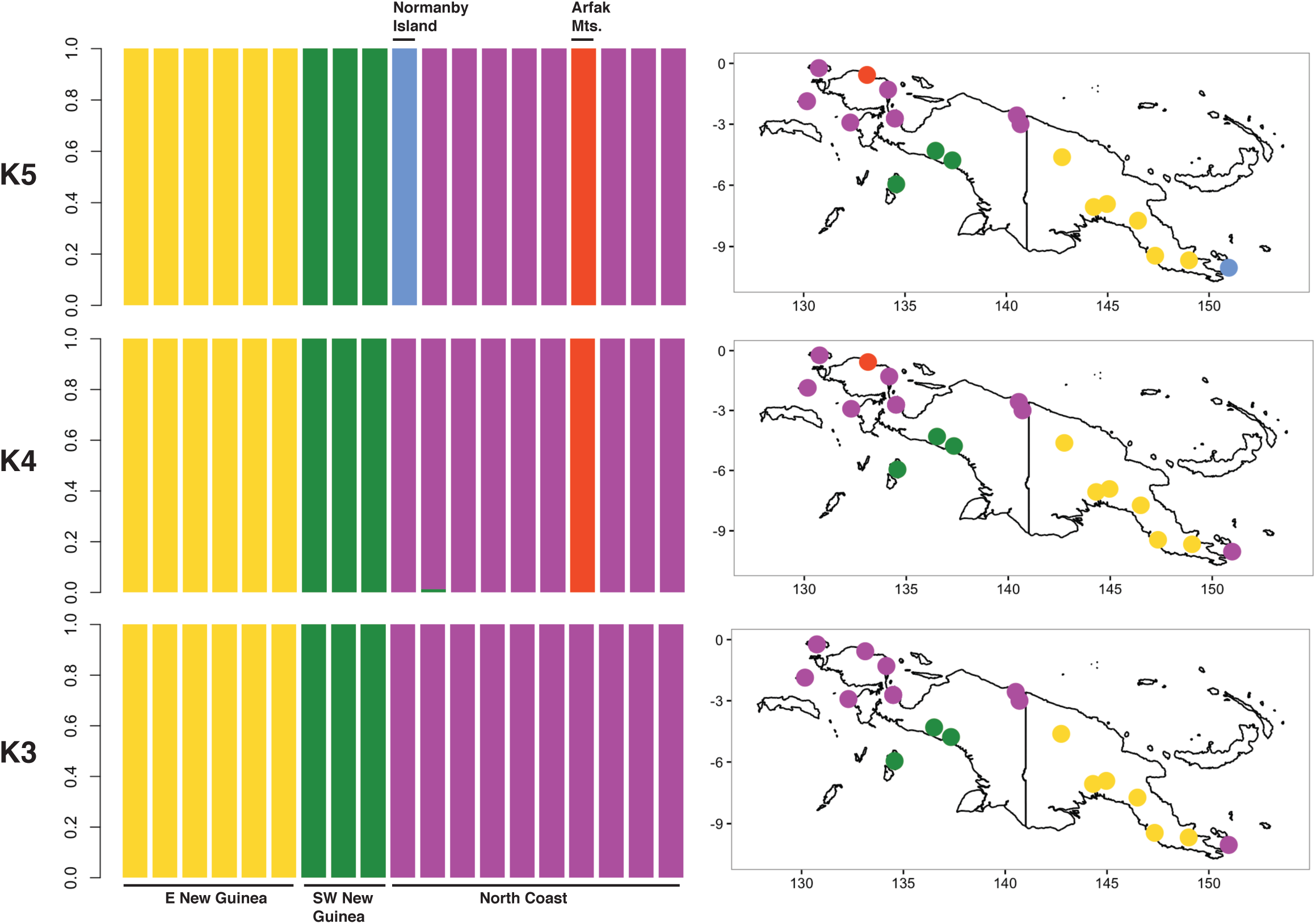
Individual membership probabilities for K=3 through K=5 ancestral populations inferred from analyzing 1,690 SNPs (100% complete data matrix) using discriminant analysis of principal components and retaining six principal component axes with two discriminant axes. Individual sampling locations are color coded accordingly.

Suchan *et al.* (2016) included historical museum specimens up to 58 years old in their validation experiment and up to 100 years old in their pilot study. We successfully captured 55 on-target loci, including four contigs exceeding 1 kb in length, from a specimen collected in 1877, and as many as 508 loci from specimens collected from 1896–1934 **(Table 1)**. Moreover, and contrary to similar analyses by Suchan *et al.* (2016) and McCormack *et al.* (2016), we found no significant relationship between age and assembly or sequencing success metrics across historical samples. While this is possibly an artifact of small sample size relative to both previous studies, it may also suggest a relatively shallow trend of degradation during a period when many bird specimens in natural history museums were originally collected. This is an encouraging result for researchers looking to make use of these specimens as a genomic resource.

### Modern DNA performs better during library preparation and sequencing, but assembly performance is similar

Our statistical analyses revealed variation between modern and ancient DNA libraries across sequencing and assembly performance metrics. Modern DNA samples had significantly higher specificity (% mapped cleaned reads), sensitivity (% of probe sequence with at least 1x coverage), and fold enrichment (the fold increase in % mapped reads over baseline random expectations) **(Figure 2)**. While initially intuitive, our findings contrast with the results of Bi *et al.* (2013) and Suchan *et al.* (2016), who found improved capture efficiency with historic samples, potentially related to smaller fragment size. We encourage future studies to explore how variation in hybridization capture protocols affect relative performance of different sample DNA sources.

Encouragingly, there were no significant differences between the two sample populations for the overall percentage of duplicate reads. However, we wish to highlight the high percentage of duplicate reads present in all samples (50.9%–91.1%). This may be due to combination of the relatively high number of amplification cycles used to amplify libraries with low input DNA prior to pooling. Nonetheless, we overcame this limitation through high sequencing effort, which allowed us to maintain sufficient coverage even after removal of duplicate reads.

Following assembly, the total number of captured on-target loci was higher among modern samples, likely reflecting higher copy number and limited degradation of DNA from fresh tissues. In contrast, mean contig length did not vary significantly among samples, indicating similar assembly performance relative to the amount of high-quality data for each sample type. While we believe this result will prove robust to different assembly methods, we encourage future studies to explore their influence on resulting assemblies and downstream analyses.

### Input DNA quantity predicts GC content, suggesting PCR bias

Our regression analyses largely failed to reveal significant correlations between input DNA / sequencing variables and assembly performance except in two comparisons **(Figure 2)**. First, an increase in the number of filtered reads was positively correlated with the total number of assembled on-target contigs, which matches a standard expectation of increased recovery with greater sampling of a genomic library with an uneven distribution of fragments, suggested for our libraries by the high levels of duplication **(Table 1)**. Second, the initial quantity of input DNA in a sample was negatively correlated with %GC content in resulting assemblies, e.g., the two samples in our study that with the lowest input DNA quantity also the highest percentage of GC content across all assembled contigs **(Table 1)**. This finding may reflect biased PCR enrichment of GC-rich exogenous microbial contamination in samples with low initial input DNA quantity (Dabney & Meyer 2012), and explain the significantly higher GC content of historic samples overall. While our SNP calling pipeline and data filtering removed sites potentially originating from non-vertebrate sequences (though see further discussion below), we nonetheless recommend researchers interested in applying hyRAD to historical specimens heed the recommendations of recent empirical studies (Gamba *et al.* 2016) to select an extraction protocol suited to degraded DNA and maximize input tissue quantity whenever possible.

### Geographic and / or taxonomic autocorrelation with input DNA type is potentially problematic with hyRAD studies

We implemented rigorous and conservative laboratory and bioinformatic protocols to reduce the influence of exogenous DNA contamination and postmortem DNA degradation, the results of which we summarize as a reference for future studies in **(Table 2)**. Despite these precautions, repeated DAPC with different parameters failed to change a basic pattern where all modern DNA samples (n=6) clustered until the chosen K value of ancestral populations was seven or more **(Table 1; Figure 5).** Unfortunately, as the close geographic proximity of modern DNA samples would also lead to this pattern, we could not easily determine from our sampling scheme if this pattern reflected biological reality or whether factors correlated with sample type, such as undetected microbial contamination, DNA degradation, and / or library amplification artifacts, were affecting population genetic inference. We first attempted to disentangle its potential drivers during PCA and DAPC analyses by examining histograms of %GC content per read for anomalous distributions, but failed to detect a significant second peak indicative of contamination with exogenous GC-rich microbes. We next performed linear regressions with specificity, sensitivity, fold enrichment, specimen age, and input DNA quantity as predictors for each of our first three retained principal components. From these regression analyses we found significant correlations of all variables with PC1, but no other PCs **(Table S1)**. While these results are consistent with the possibility that biased sequencing performance affected population genetic inference, the exact mechanism responsible remains unclear. We suggest researchers intending to use hyRAD in studies with both modern and historic tissue attempt to avoid geographic and taxonomic autocorrelation wherever possible, and include a control in their sampling scheme to indicate potential DNA input quality problems.

### The phylogeography of S. torotoro reflects biogeographic barriers in New Guinea

Our DAPC results are consistent with previous studies of lowland avian phylogeography in New

Guinea, and we interpret them as independent confirmation the ability of hyRAD to reveal biologically meaningful patterns. K-means clusters recovered for three to five hypothetical ancestral populations reflect broad trends in codistributed taxa and expected patterns of genetic variation given geographic barriers and the geologic history of New Guinea **(Figure 5)** (Dumbacher & Fleischer 2001; Deiner *et al.* 2011). Although IBD across all samples was not significant (see *Results* for details), the initial division of samples into mainland eastern and western clusters is consistent with both the primary latitudinal axis of the island and the barriers to gene flow in lowland forest taxa presented by the Bewani Mountains and Trans-Fly savannah region (Dumbacher & Mack 2007; Deiner *et al.* 2011). Though counter intuitive, inclusion of Normanby Island subspecies *S. t. ochracea* in the western New Guinea cluster is consistent with previous studies that have reported genetic similarity between other taxa in far eastern and far western New Guinea, such as birds of the genus *Pitohui* (Dumbacher & Fleischer 2001).

When K=3, samples from southwest New Guinea clustered with those from the Aru Islands, suggesting shared ancestry among these currently allopatric populations. This is potentially explained by both the linkage of these landmasses during the Pleistocene via the Sahul Shelf (Voris 2000) and the subsequent emergence of previously identified barriers to avian gene flow to the north, east, and west in the form of the Central Ranges, the Trans-Fly Savannah, and Aetna Bay, respectively (Dumbacher & Fleischer 2001; Deiner *et al.* 2011). K=4 added a cluster containing of a single individual from the northern slope of the Arfak mountains, which may indicate a previously undiagnosed biogeographic unit in the Vogelkop Peninsula. However, as the sample in question was an outlier in age and assembly size **(Table 1)**, we are reluctant to over interpret this signal. Regardless, our analysis reveals broad similarities between the phylogeography of *Syma torotoro* and the codistributed lowland bird species *Colluricincla megarhyncha* (Deiner et al. 2011), albeit with lower resolution due to the inherent limitations of our sampling. We believe that future studies of resident lowland forest species with similar ranges that use hyRAD or other means of capturing nuclear DNA markers will continue to aid in building a cohesive picture of the comparative phylogeography of this biodiverse region.

## ACKNOWLEDGEMENTS

For tissue loans we thank R. Moyle and M. Robbins at KUMNH, P. Sweet at AMNH, and Robert Prys-Jones at NHMUK-Tring. For field assistance, we thank T. Bozic, A. Mack, D. Mindell, J. Fuchs, M.A.C. Dodge, and Bruno and Carmen Montel. For laboratory assistance, we thank L. Wilkinson and M. Flannery. Support for this research was provided by NDSEG and WRF-Hall Fellowships, a Society of Systematic Biology GRSA, and a gift from M. and K. Simon. For comments on the manuscript, we thank J. Klicka, C.J. Battery, D. Slager, C. French, and R. Harris. We performed all laboratory work at the Center for Comparative Genomics, California Academy of Sciences, and all sequencing at the Vincent J. Coates Genomics Sequencing Laboratory at UC Berkeley, supported by NIH Instrumentation Grant S10 0D018174. In PNG, we thank the Institute of Biological Research, the National Museum and Art Gallery, the National Research Institute, and the Department of Conservation. We add special thanks to the many New Guineans who provided field and logistical assistance, especially Bulisa Iova and tribal leaders of the villages that participated in and allowed our work.

